# Activation of Rab11a and endocytosis by Phosphatidylinositol 4-kinase III beta promotes oncogenic signaling in breast cancer

**DOI:** 10.1101/2020.02.10.942177

**Authors:** Spencer A MacDonald, Katherine Harding, Patricia Bilodeau, Christiano T de Souza, Carlo Cosimo Campa, Emilio Hirsch, Rebecca C Auer, Jonathan M Lee

## Abstract

Endosomes are now recognized as important sites for regulating signal transduction. Here we show that the lipid kinase phosphatidylinositol 4-kinase III beta (PI4KIIIβ) regulates both endocytic kinetics and receptor signaling in breast cancer cells. PI4KIIIβ generates phosphatidylinositol 4-phosphate from phosphatidylinositol and is highly expressed in a subset of breast cancers. However, the molecular mechanism by which PI4KIIIβ promotes breast cancer is unclear. We demonstrate that ectopic PI4KIIIβ expression increases the rates of both endocytic internalization and recycling. PI4KIIIβ deletion reduces endocytic kinetics accompanied by a concomitant decrease in activity of the Rab11a GTPase, a protein required for endocytic function. Finally, we find that PI4KIIIβ activates IGF-IRβ signaling dependent on endosome function. Regulation of endocytic function by PI4KIIIβ is independent of its kinase activity but requires interaction with the Rab11a. This suggests that PI4KIIIβ controls endosomal kinetics and signaling by directly modulating Rab11a function. Our work suggests a novel regulatory role for PI4KIIIβ in endosome function and plasma membrane receptor signaling.

## INTRODUCTION

Phosphatidylinositol 4-kinase III beta (PI4KIIIβ) generates phosphatidylinositol 4-phosphate (PI4P) from phosphatidylinositol (PI) and the protein has a well-characterized role in the maintenance of the structure and function of the Golgi^2, 3, 10^. Recent evidence suggests that PI4KIIIβ has a direct role in the progression and development of cancer^43^. For example, transcriptional analysis of 1992 primary human breast tumours by Curtis *et al.* identified *PI4KB*, the gene encoding human PI4KIIIβ, as a cancer driver gene due to its frequent gene amplification and overexpression in primary tumour tissue^7^. Additionally, we have reported that the PI4KIIIβ protein is highly expressed in ∼20% of primary human breast tumours^28^. Consistent with a functional role for PI4KIIIβ in cancer, ectopic PI4KIIIβ expression also disrupts three-dimensional epithelial morphogenesis of breast cells and promotes cell motility and actin remodeling^1, 20, 31^. Finally, our lab has reported that PI4KIIIβ is able to directly activate Akt signaling ^28^.

PI4KIIIβ has kinase-independent functions and could therefore regulate cancer progression through mechanisms distinct from PI4P generation. In *Drosophila*, for example, the PI4KIIIβ homolog four-wheel drive (*Fwd*) regulates Rab11localization and endocytic function independent of PI4P^32^. Rab11a is a member of the Rab family of small GTPases that control membrane identity and function in the endocytic system^39^. PI4KIIIβ directly binds Rab11^5^ and *Fwd*/Rab11interaction is necessary for normal cytokinesis in *Drosophila* spermatocytes and loss of *Fwd* leads to sterility in males ^32^. In yeast and mammalian cells, PI4KIIIβ is similarly necessary for the binding and localization of Rab11a to the Golgi independent of PI4P^9, 32^. Additionally, PI4KIIIβ-mediated activation of Akt requires Rab11a interaction but not an active lipid kinase domain^28^. Taken together, PI4KIIIβ appears to have an important functional relationship with Rab11a and endosome function independent of its ability to generate PI4P.

The endocytic system is best known for moving cargo to and from the plasma membrane ^23^. In addition, by controlling both plasma membrane abundance and degradation of signalling receptors, the endocytic system controls both activation, propagation and timing of signalling cascades downstream tumorigenic receptors^37, 38, 40^. For example, in endosomes the ligand-bound epidermal growth factor receptor (EGFR) and the small GTPase Rab5 recruit the adaptor protein APPL promoting the activation of Akt^24, 35^. We have previously shown that the interaction between PI4KIIIβ and Rab11a is required for activation of Akt signaling^28^. We therefore hypothesized that the interaction between PI4KIIIβ and endosomes is related to its role PI4KIIIβ in oncogenesis. Here we find that PI4KIIIβ regulates both endocytosis and recycling of surface receptors affecting Rab11a activation. Our findings suggest that PI4KIIIβ has an important role in endosome function and endosome-mediated signaling in cancer cells.

## RESULTS

### PI4KIIIβ regulates mammary tumourigenesis

We have previously shown that PI4KIIIβ protein is highly expressed in a ∼20% fraction of human breast cancers^28^. In order to determine whether or not PI4KIIIβ might have a role in primary mammary tumour development, we used CRISPR/Cas9 to delete *Pi4kb*, the mouse PI4KIIIβ homolog, from the mouse mammary 4T1 tumour cell line (Figure 1). 4T1 is a metastatic derivative of a spontaneous mammary tumour from Balb/c mice ^33^. When injected into the mammary fat pad of syngeneic mice they grow as solid tumours and this model is commonly used to study *in vivo* mammary tumour growth^26^. Two independent lines of 4T1 that lack *Pi4kb* grow substantially slower than control cells (Figures 1a-d). Re-expression of human PI4KIIIβ in the *Pi4kb* null cells returns tumorigenicity to wildtype levels (Figure 1e). These results are consistent with the idea that PI4KIIIβ function has an important functional role in mammary tumour development.

**Figure 1.**
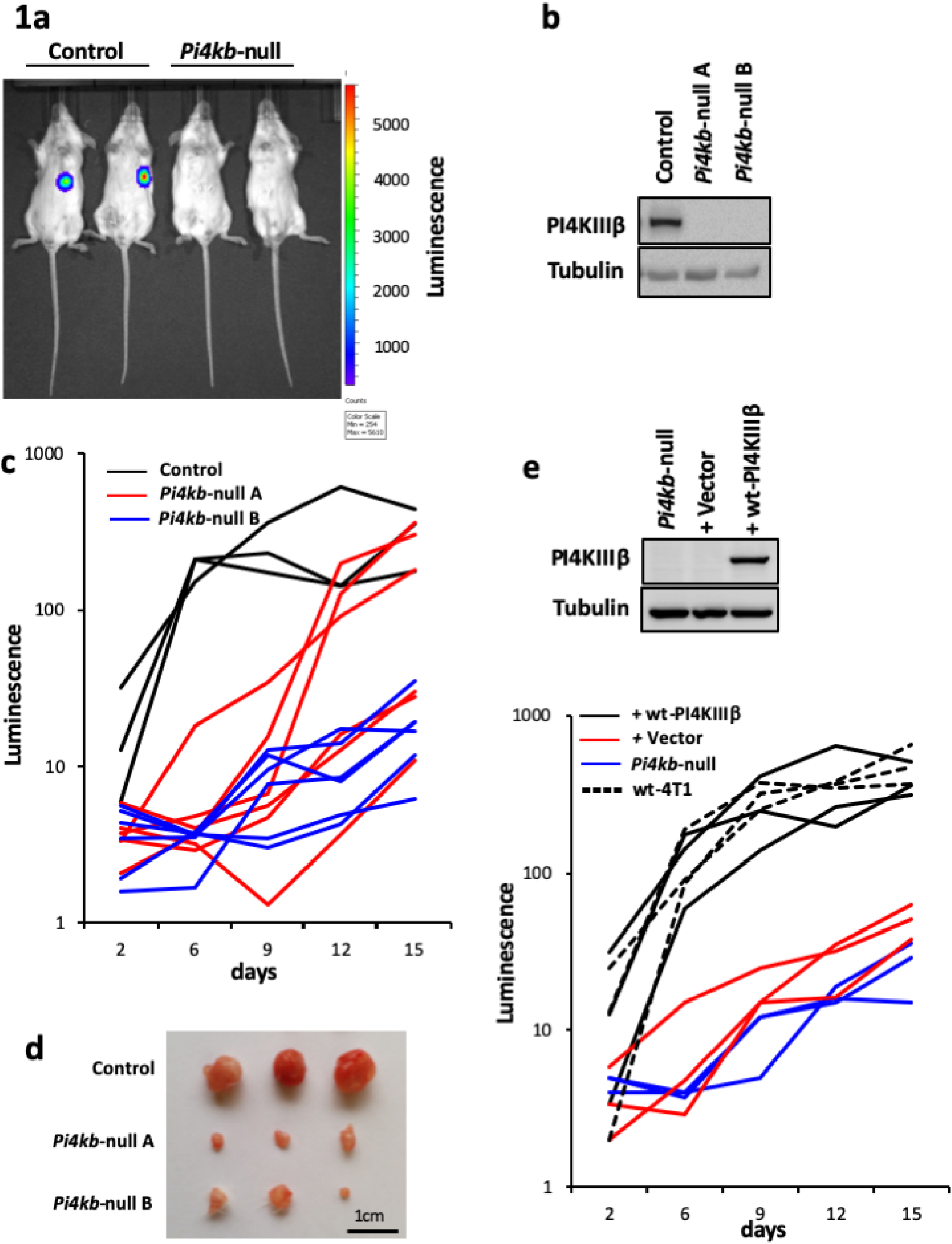
*Pi4kb* deletion inhibits mammary tumorigenesis in mice. (**a**) Representative image of tumour formation in Balb/c mice injected with wildtype 4T1 cells (Control) and mice injected with *Pi4kb* deleted 4T1 cells (*Pi4kb*-null). The tumours shown are from day 7 post injection. Tumour volume is indicated colorimetrically. (**b**) Western blot analysis showing levels of PI4KIIIβ (110kDa) with Tubulin (55kDa) as a loading control in 4T1 Control and two independent CRISPR/Cas9 knockouts of *Pi4kb* gene (*Pi4kb*-null A and B) cell lines. (**c**) Tumour development as a function of time in 4T1 Control (black line), and *Pi4kb*-null A and B (red and blue lines) cell lines. Mice were euthanized at day 15 post injection. The data shown represents 4 independent experiments. (**d**) Representative tumours from Control and *Pi4kb*-null A and B injected 4T1 cell lines. Tumours are from day 9 post injection. (**e**) *Upper Panel.* Expression of PI4KIIIβ in *Pi4kb*- null 4T1 cells and those rescued by expression of wildtype human *PI4KB* or empty vector. *Lower panel.* Tumour development as a function of time in *Pi4kb*-null cells (blue), *Pi4kb*-null cells rescued by human *PI4KB* (solid black), human *PI4KB* with empty vector (red) and wildtype 4T1 cells (hatched black). Mice were euthanized at day 15 post injection. The data shown represents 2 independent experiments.

### PI4KIIIβ activates IGF-IRβ

We hypothesized that PI4KIIIβ is oncogenic through an ability to activate receptor signaling. We have previously reported that ectopic expression of PI4KIIIβ activates Akt in a kinase-independent manner in breast cell lines^28^. To determine whether or not PI4KIIIβ affects signaling pathways in addition to Akt, we expressed ectopic wildtype or kinase-inactive (D656A) PI4KIIIβ in BT549 human breast ductal carcinoma cells (Figure 2a). We observed a significant 2-3-fold increase in IGF-IRβ activation when either wild-type or kinase-inactive PI4KIIIβ was highly expressed compared to controls (Figures 2b and c). This demonstrates that PI4KIIIβ activates in IGF-IRβ signaling in a manner that does not require an active lipid kinase domain.

**Figure 2.**
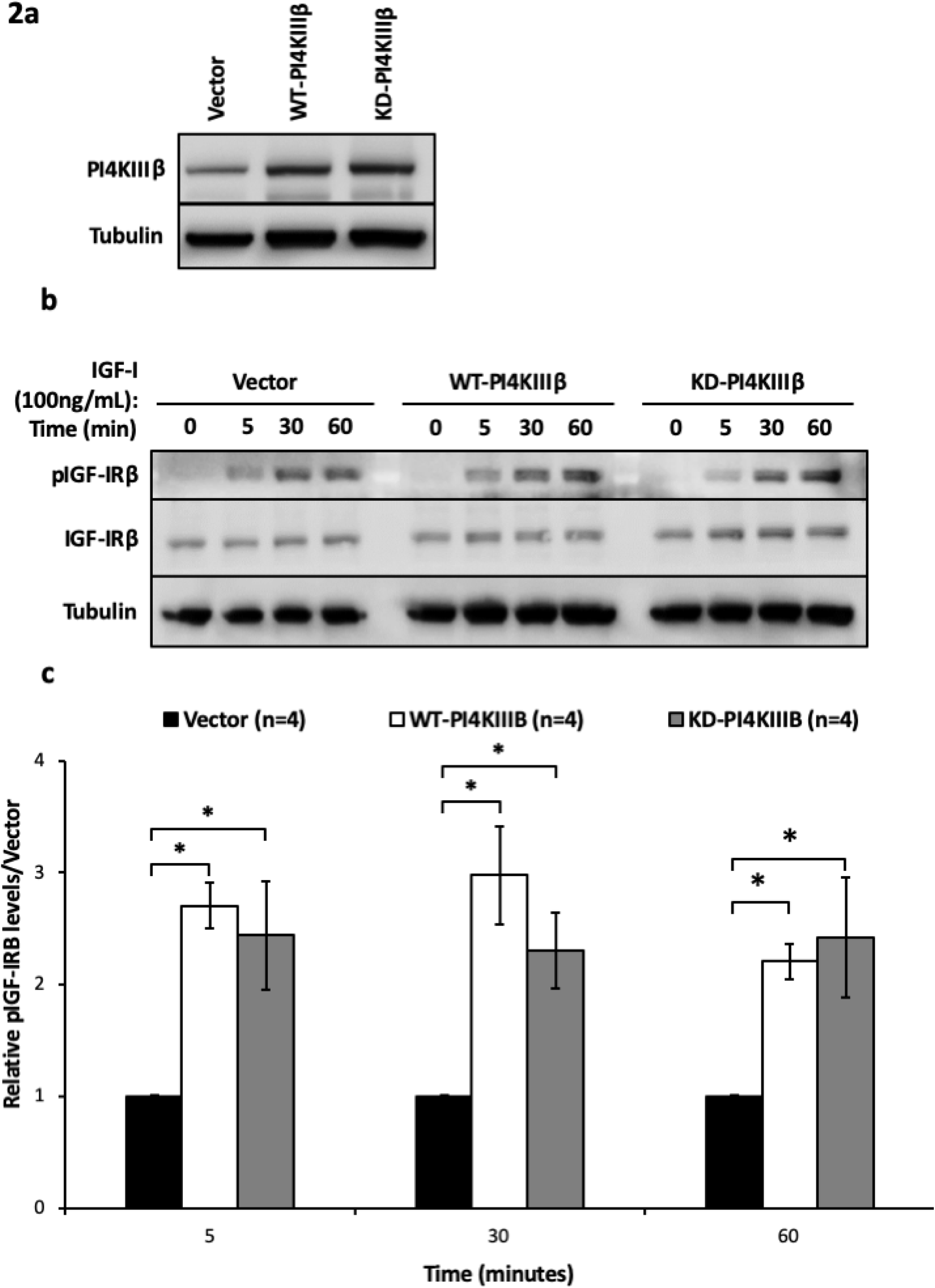
PI4KIIIβ activates IGF-IRβ signaling. (**a**) Western blot analysis showing levels of PI4KIIIβ (110kDa) with Tubulin (55kDa) as a loading control in BT549 vector control (Vector), wildtype PI4KIIIβ-overexpressing (WT-PI4KIIIβ), and kinase-inactive PI4KIIIβ-expressing (KD-PI4KIIIβ) human breast ductal carcinoma cell lines. Each lane contains 30μg of total protein. (**b**) BT549 Vector, WT-PI4KIIIβ, and KD-PI4KIIIβ cell lines were serum starved overnight followed by stimulation with IGF-I (100ng/mL) for the indicated time periods. Lysate was collected and subjected to western blot analysis to determine levels of pIGF-IRβ (95kDa) and IGF-IRβ (95kDa) with Tubulin (55kDa) as a loading control. Each lane contains 30µg of total protein. (**c**) Protein levels were quantified by densitometry and the data shown represents the mean ± SE of the mean from 4 independent trials comparing the pIGF-IRβ levels relative to the IGF-IRβ levels in each cell type, followed by normalization to the Vector for each time point.

### PI4KIIIβ expression alters endocytic kinetics

The activation of Akt by PI4KIIIβ previously established by our lab was shown to be dependent upon the presence of Rab11a^28^. Because Rab11a regulates endosome recycling, we hypothesized that the effects of PI4KIIIβ on breast cancer oncogenesis may be due to a regulatory role in endosome function^23^. To determine whether PI4KIIIβ might affect transport of surface receptors from endosomes to plasma membrane, we used transferrin pulse-chase assays to measure the rate of endocytic recycling. We observed that BT549 cells overexpressing wildtype or kinase-inactive PI4KIIIβ were able to recycle out twice as much internalized transferrin as the vector control (Figures 3a and b). Transferrin receptor expression is similar between the cell lines (Supplementary Figure 1) and recycling kinetics are normalized for each cell line to account for small differences in initial transferrin binding to the cell surface. Next, we used transferrin uptake assays to measure the rate at which cells were able to internalize fluorescent transferrin. BT549 cells with ectopic wildtype or kinase-inactive PI4KIIIβ were able to internalize significantly more transferrin (4-5x) than the vector control (Figures 3c and d). Uptake kinetics were normalized for each cell line to account for any difference in initial transferrin binding to the cell surface.

**Figure 3.**
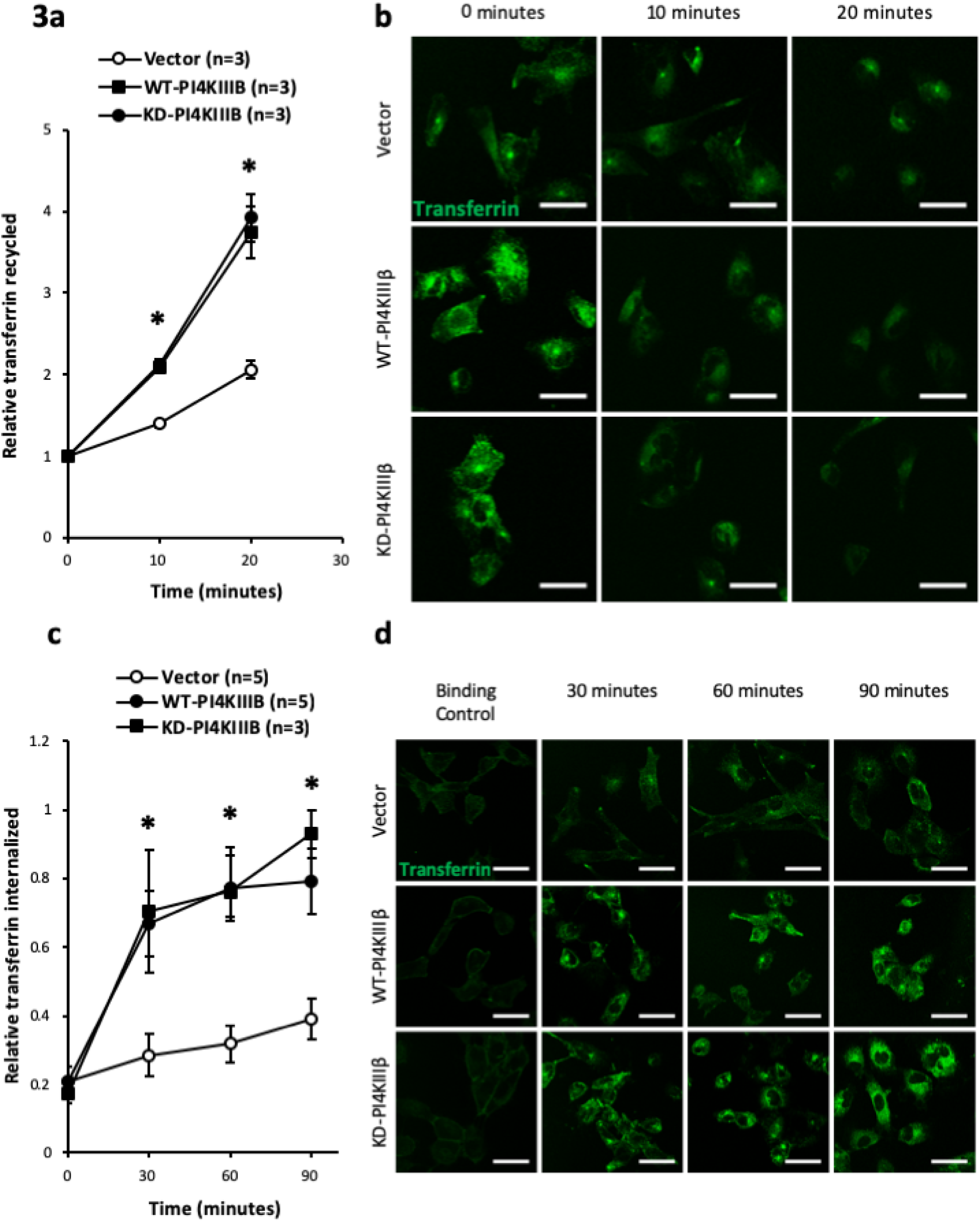
High PI4KIIIβ expression increases rates of endocytic activity. (**a**) The relative transferrin recycled in BT549 vector control (Vector), wildtype PI4KIIIβ-overexpressing (WT-PI4KIIIβ), and kinase-inactive PI4KIIIβ-expressing (KD-PI4KIIIβ) human breast ductal carcinoma cell lines was calculated by determining the average corrected total cellular fluorescence for 25 cells in each condition per trial. The relative transferrin recycled is presented as the mean ± SE of the mean, from 3 independent trials, following background subtraction and normalization to the initial time point for each cell line. Statistical significance (*, *P* < 0.05, one-way ANOVA with multiple comparison tests) is indicated. (**b**, **d**) Representative confocal images of paraformaldehyde fixed BT549 Vector, WT-PI4KIIIβ, and KD-PI4KIIIβ cell lines after undergoing transferrin (**b**) pulse-chase or (**d**) uptake assay for specified times. 63x magnification. Scale bars, 50µm. (**c**) The relative transferrin internalized was calculated by determining the average corrected total cellular fluorescence for 25 cells in each condition per trial. The relative transferrin internalized is presented as the mean ± SE of the mean, from 5 (Vector, KD-PI4KIIIβ) or 3 (WT-PI4KIIIβ) independent trials, following background subtraction and normalization to the binding control for each cell line. Statistical significance (*, *P* < 0.05, one-way ANOVA with multiple comparison tests) is indicated.

To further explore the importance of PI4KIIIβ on endosome function, we created *PI4KB*-null cell lines using CRISPR/Cas9 in BT549 wildtype cells (Figure 4a). *PI4KB*-null cells are viable and appear to proliferate normally. Perhaps surprisingly, we did not observe any disruption of the gross morphology of either the cis- or trans-Golgi in *PI4KB*-null cells (Figures 4b and c). In order to investigate how the deletion of PI4KIIIβ might affect endocytic kinetics, we repeated the transferrin recycling and uptake assays in *PI4KB*-null cell lines. We found that BT549 cells with *PI4KB* deletion were able to recycle only half as much transferrin as the wildtype cells (Figures 4d and e). Furthermore, cells with PI4KIIIβ deletion internalized significantly less transferrin (4-5x) than the wildtype (Figures 4f and g). Uptake and recycling kinetics were normalized for each cell line to account for any difference in initial transferrin binding or internalization to the cell surface. The ability of PI4KIIIβ to regulate endosome function is not the result of changes in recycling endosome size or number. We used immunofluorescence to measure the size and number of Rab11a-positive vesicles and found that there was no detectable difference in the amount or size of Rab11a vesicles between wildtype cells and those with no or high PI4KIIIβ expression (Supplementary Figure S2). Overall, our results are consistent with a novel role for PI4KIIIβ in controlling endocytic internalization and recycling.

**Figure 4.**
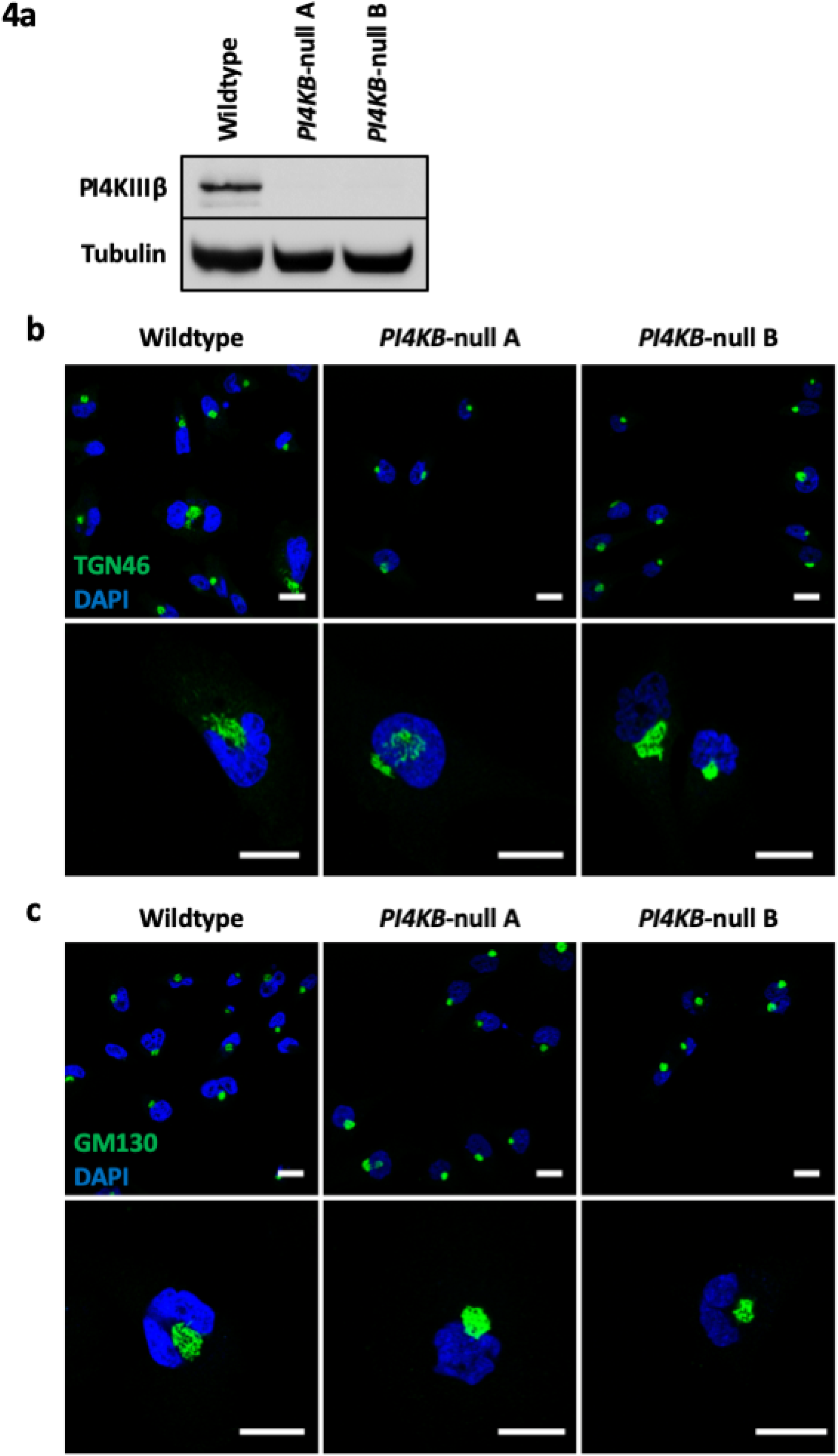
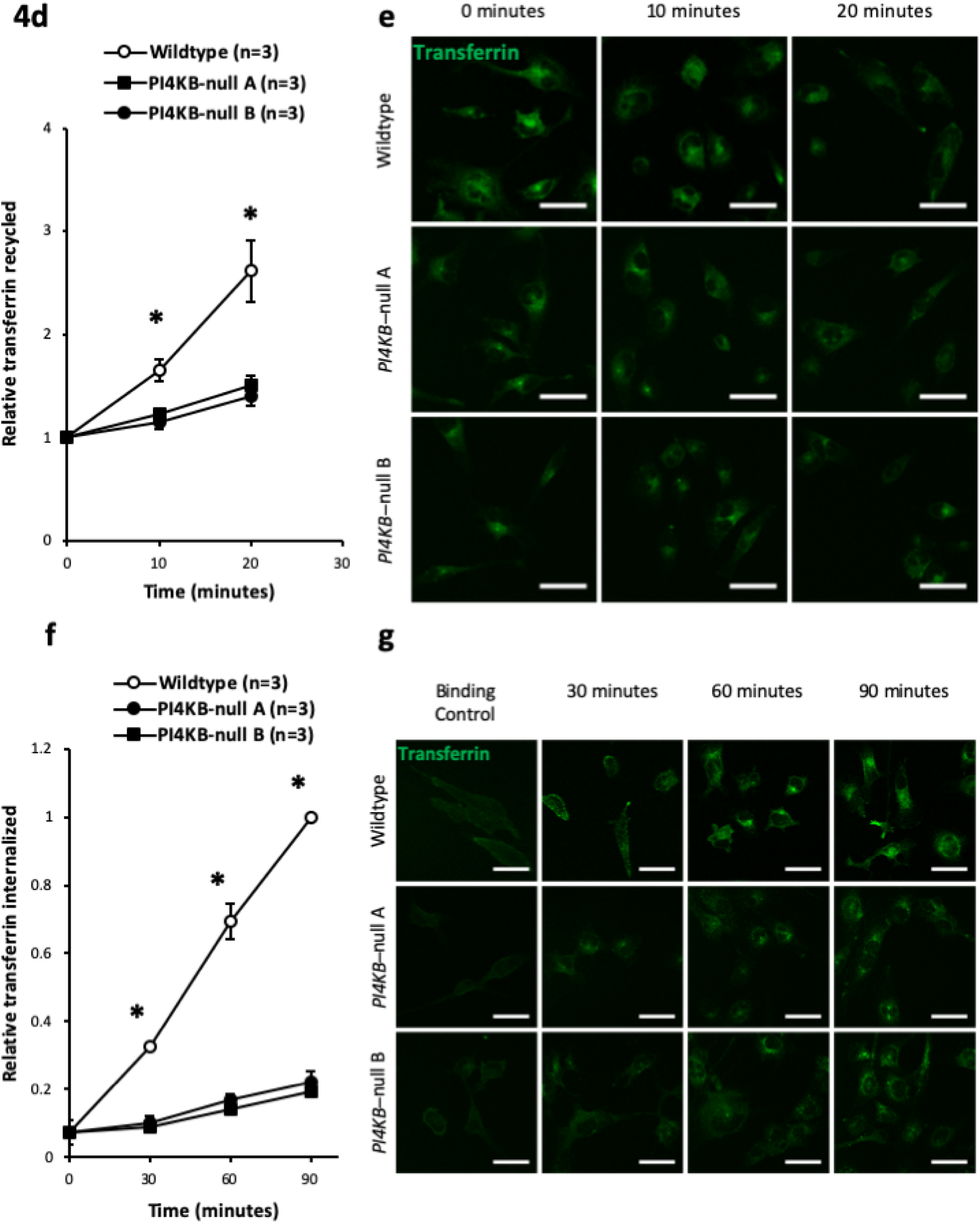
Deletion of PI4KIIIβ decreases endocytic activity rates. (**a**) Western blot analysis showing levels of PI4KIIIβ (110kDa) with Tubulin (55kDa) as a loading control in BT549 wildtype (Wildtype) and two independent CRISPR/Cas9 knockouts of PI4KIIIβ (*PI4KB*-null A and B) human breast ductal carcinoma cell lines. Each lane contains 30µg of total protein. (**b**, **c**) Representative confocal images of paraformaldehyde fixed BT549 Wildtype, and *PI4KB*-null A and B cell lines following immunofluorescence of (**b**) *trans-*Golgi marker, TGN46 and (**c**) *cis*-Golgi marker, GM130. 63x magnification. Scale bars, 20µm. (**d**) The relative transferrin recycled was calculated by determining the average corrected total cellular fluorescence for 25 cells in each condition per trial. The relative transferrin recycled is presented as the mean ± SE of the mean, from 3 independent trials, following background subtraction and normalization to the initial time point for each cell line. Statistical significance (*, *P* < 0.05, one-way ANOVA with multiple comparison tests) is indicated. (**d**, **g**) Representative confocal images of paraformaldehyde fixed BT549 Wildtype, and *PI4KB*-null A and B cell lines after undergoing transferrin (**d**) pulse-chase or (**g**) uptake assay for specified times. 63x magnification. Scale bars, 50µm. (**f**) The relative transferrin internalized was calculated by determining the average corrected total cellular fluorescence for 25 cells in each condition per trial. The relative transferrin internalized is presented as the mean ± SE of the mean, from 3 independent trials, following background subtraction and normalization to the binding control cell line. Statistical significance (*, *P* < 0.05, one-way ANOVA with multiple comparison tests) is indicated.

### Lipid kinase-independent control of endocytic function

PI4KIIIβ is a multifunctional protein and we next wanted to determine which of the documented roles of PI4KIIIβ are involved in regulating endocytic function. To this end, we expressed wild-type, kinase-inactive (D656A) and Rab11a-binding deficient (N162A) PI4KIIIβ in the BT549 *PI4KB*-null CRISPR/Cas9 cell lines (Figure 5a). We then used the transferrin pulse-chase and uptake assays to measure the efficiency of both receptor internalization and recycling. We found that the reintroduction of wildtype and kinase-inactive PI4KIIIβ was able to rescue the rate of transferrin recycling (Figures 5b and c). However, the Rab11a-binding deficient PI4KIIIβ was unable to do so (Figures 5b and c). This is consistent with the idea that endocytic recycling by PI4KIIIβ does not require lipid kinase activity but is dependent upon the functional interaction between PI4KIIIβ and Rab11a.

**Figure 5.**
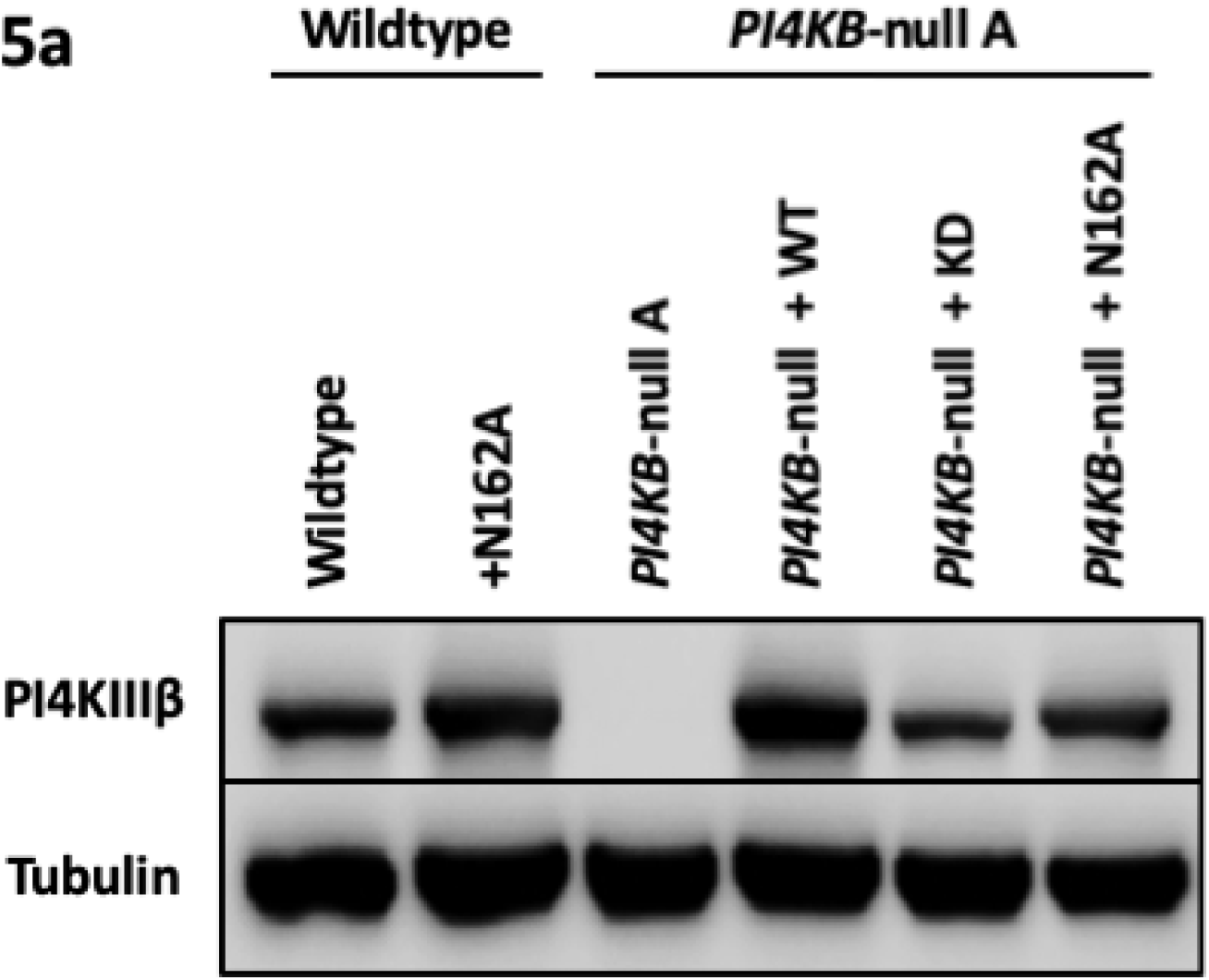
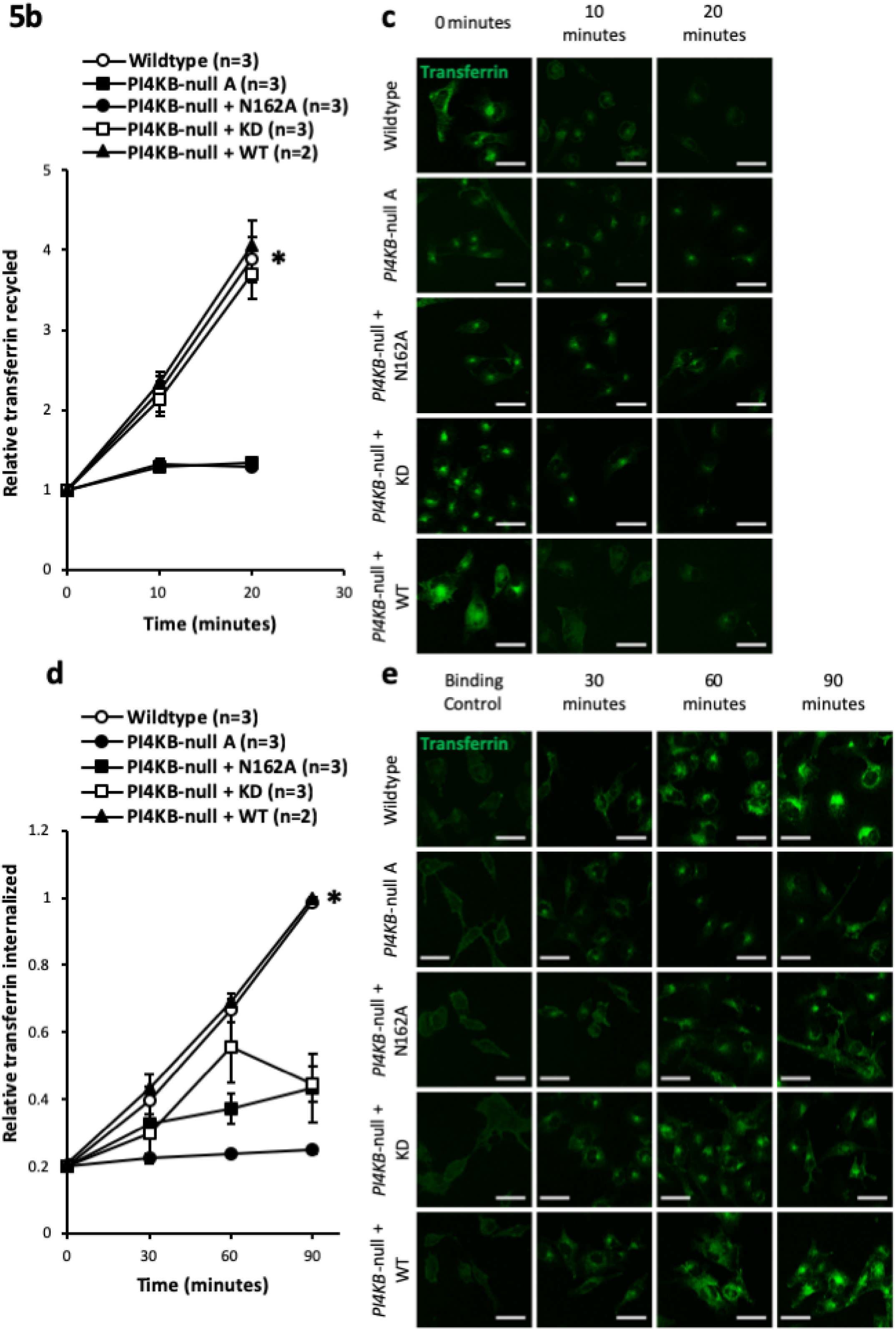
Reintroduction of PI4KIIIβ rescues recycling and internalization of transferrin. (**a**) Western blot analysis showing levels of PI4KIIIβ (110kDa) with Tubulin (55kDa) as a loading control in BT549 wildtype cell line transfected with PI4KIIIβ unable to bind Rab11a (+N162A), and *PI4KB*-null A cell line transfected with wildtype PI4KIIIβ (*PI4KB*-null + WT), kinase-inactive PI4KIIIβ (*PI4KB*-null + KD), or PI4KIIIβ unable to bind Rab11a (*PI4KB*-null + N162A) plasmids followed by antibiotic selection in order to generate stable cell lines. Each lane contains 30µg of total protein. (**b**) The relative transferrin recycled was calculated by determining the average corrected total cellular fluorescence for 25 cells in each condition per trial. The relative transferrin recycled is presented as the mean ± SE of the mean, from 3 (Wildtype, *PI4KB*-null A, *PI4KB*-null + N162A, *PI4KB*-null +KD) or 2 (*PI4KB*-null + WT) independent trials, following background subtraction and normalization to the initial time point for each cell line. Statistical significance (*, *P* < 0.05, one-way ANOVA with multiple comparison tests) is indicated. (**c**, **e**) Representative confocal images of paraformaldehyde fixed BT549 Wildtype, *PI4KB*-null A, *PI4KB*-null + N162A, *PI4KB*-null + KD, and *PI4KB*-null + WT cell lines after undergoing transferrin (**c**) pulse-chase or (**e**) uptake assay for specified time. 63x magnification. Scale bars, 50µm. (**d**) The relative transferrin internalized was calculated by determining the average corrected total cellular fluorescence for 25 cells in each condition per trial. The relative transferrin internalized is presented as the mean ± SE of the mean, from 3 (Wildtype, *PI4KB*-null A, *PI4KB*-null + N162A, *PI4KB*-null + KD) or 2 (*PI4KB*-null + WT) independent trials, following background subtraction and normalization to the binding control for each cell line. Statistical significance (*, *P* < 0.05, one-way ANOVA with multiple comparison tests) is indicated.

On the other hand, we found that the rate of transferrin uptake was only fully rescued if the reintroduced PI4KIIIβ was catalytically active and able to interact with Rab11a (Figures 5d and e). In the absence of endogenous PI4KIIIβ, kinase-inactive or Rab11a-binding deficient PI4KIIIβ resulted in only a partial (∼30-50%) rescue of the transferrin uptake rate (Figures 5d and e). This is in contrast to those obtained with recycling, where we observed a similar increase in recycling between wildtype and kinase-inactive PI4KIIIβ (Figures 5b and c). This could be due to the lower levels of kinase-inactive PI4KIIIβ expression in our *PI4KB*-null rescue cell lines compared to the amount of wildtype PI4KIIIβ (Figure 5a). However, this lower level of kinase-inactive PI4KIIIβ expression fully rescued recycling. We propose that the kinase-dependent and -independent functions of PI4KIIIβ may cooperate in regulating endosome internalization.

### PI4KIIIβ is found in some endosomal compartments

We and others have previously reported that ectopically expressed PI4KIIIβ, both wildtype and kinase-inactive, colocalizes with Rab11a in recycling endosome^28^. Purified PI4KIIIβ and Rab11a proteins bind each other with 1:1 stoichiometry^5^. Since we have found that PI4KIIIβ has an impact on both endocytic recycling and internalization, we speculated that PI4KIIIβ might be found in endocytic compartments. To test this idea, we used immunofluorescence to determine whether or not PI4KIIIβ could be found in early or late endosomes. As shown in Figure 6, PI4KIIIβ can be found in early endosomes, as identified by those with EEA1 and Rab5 staining, as well as in late endosomes, i.e. those staining with the late endosome marker Rab7. The presence of PI4KIIIβ in these endosomes occurs in both wildtype BT549 cells (Figures 6a-c) and those ectopically expressing wildtype or kinase-inactive PI4KIIIβ and the vector control (Figure 6d). Coupled with our kinetic data on receptor internalization (Figures 4 and 5), the presence of PI4KIIIβ in a subset of early, late, and recycling endosomes further indicates that PI4KIIIβ regulates endosome functions in addition to that of Rab11a-dependent recycling.

**Figure 6.**
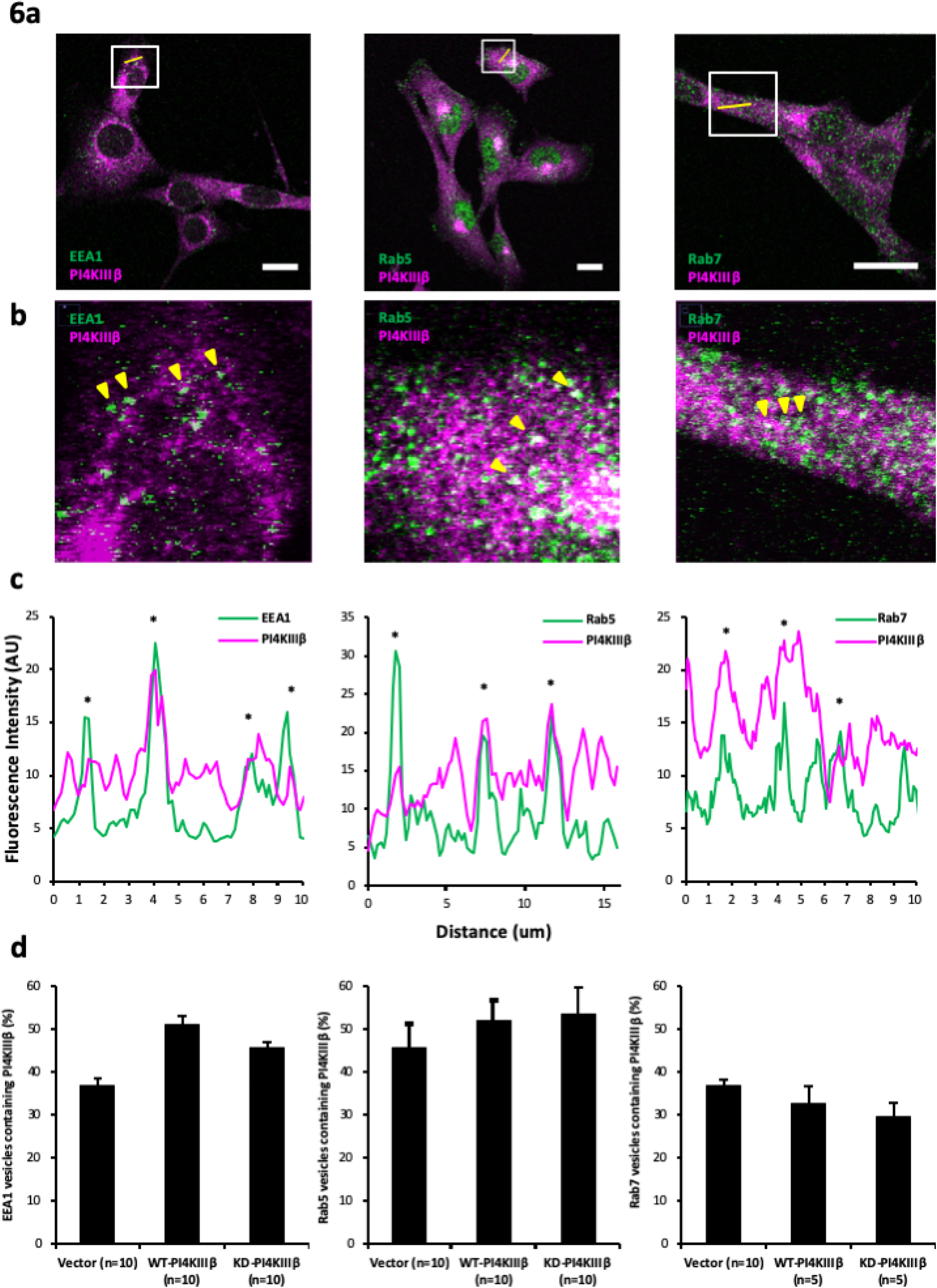
PI4KIIIβ is found in early and late endosomes. (**a**) Representative confocal images of EEA1, Rab5, Rab7 (left to right) (green), and PI4KIIIβ (magenta) in paraformaldehyde fixed BT549 wildtype PI4KIIIβ-overexpressing (WT-PI4KIIIβ) cells. 63x magnification. Scale bars, 20µm. (**b**) Zoomed in image of white insets in Figure 5a with coincident fluorescence marked (yellow arrows). (**c**) Intensity profile for EEA1, Rab5, Rab7 (left to right) (green), and PI4KIIIβ (magenta) across the yellow line drawn in Figure 5a. Coincident fluorescence (*) is indicated. (**d**) The percentage of EEA1, Rab5, or Rab7 (left to right) vesicles that have coincident fluorescence with PI4KIIIβ in each of the BT549 vector control (Vector), WT-PI4KIIIβ, and kinase-inactive PI4KIIIβ expressing (KD-PI4KIIIβ) cell lines. At least 50 EEA1, Rab5, or Rab7 vesicles were counted per cell and the number of cells counted for each type is shown.

### PI4KIIIβ regulates Rab11 activity

To further explore a role for PI4KIIIβ in endosome function, we next explored whether or not Rab11a activity might be depend on PI4KIIIβ. We used a FRET (Fluorescence Resonance Energy Transfer)-based Rab11a reporter^6^ to measure Rab11a activity in wild-type and CRISPR deleted BT549 cells (AS-Rab11). This reporter consists of human Rab11a, the C-terminal region of the FIP3 Rab11a binding protein^15^, a modified monomeric yellow fluorescent protein (mcpVenus), monomeric cyan fluorescent protein (mECFP), and human Rab11a (Fig 7a). In this reporter, activation of Rab11a promotes interaction between the FIP3 C-terminal region and Rab11a, increasing FRET because of a change in the orientation of the Venus and CFP fluorophores. FRET is represented by the 525nm/475nm (FRET/CFP) emission ratio^27, 30^ (Fig. 7a). With the AS-Rab11 biosensor, wild type BT549 cells show Rab11 activity in the perinuclear region and in some peripheral vesicles. These regions are likely to be the Golgi and endocytic vesicles respectively. CRISPR deletion of PI4KIIIβ in BT549 cells reduces the overall amount of Rab11 activity per cell (Fig 7b). Wild-type BT549 cells have, on average, a FRET 525nm/475nm ratio of 3.59 ±0.6, significantly (p<0.000063, *t-test*) greater than the 3.03 ±0.6 average in CRISPR cells. The FRET signal in the CRISPR cells is statistically indistinguishable from the FRET signal in cells expressing the AS-Rab11^S25N^ negative control (3.25 ±0.6). In AS-Rab11^S25N^, wild-type Rab11a has been replaced with a dominant-negative mutant. Our observation that loss of PI4KIIIβ causes a decrease in Rab11a activity is consistent with the idea that PI4KIIIβ is a direct activator of Rab11a.

**Figure 7.**
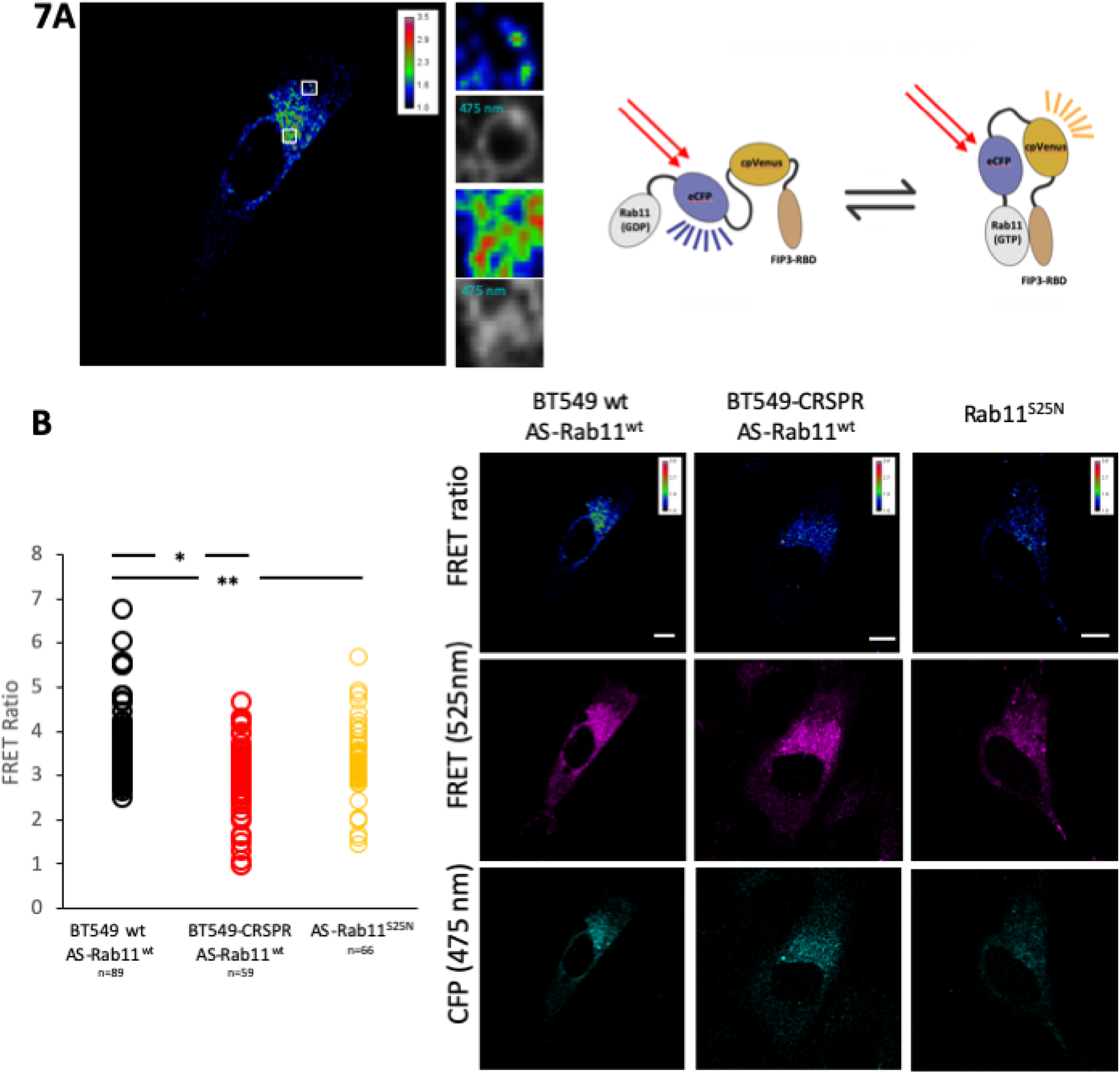
Loss of PI4KIIIβ causes a decrease in cellular Rab11a activity. **A)** *Left panel*. A FRET reporter was used to measure Rab11a activity in BT549 cells. Rab11a activity is measured as the ratio of the 525nm/475nm fluorescence and is depicted colorimetrically. Boxes show enlarged portions of the FRET ratio along with the levels of the reporter depicted by a monochrome display of the 475nm channel. *Right Panel.* Schematic of the domain structure of the FRET reporter used. **B)** *Left Panel.* graph shows mean the mean FRET ratio of wild-type BT549 cells (n=89), CRSPR deleted BT549 cells (n=59), and control cells (n=66. CRSPR deleted BT549 cells) expressing the dominant negative AS-Rab11^S25N^ mutant. Statistical significance (t-test) is indicated (* p<0.000063). *Right Panels*. Representative images of cells used for quantitation. Rab11a activity is measured as the ratio of the 525nm/475nm fluorescence and is depicted colorimetrically. Boxes show enlarged portions of the FRET ratio along with the levels of the reporter depicted by a monochrome display of the 475nm channel.

### PI4KIIIβ mediated activation of IGF-IRβ is dependent on endosome function

Because we saw a significant increase in the activation of IGF-IRβ in cells with high PI4KIIIβ expression (Figures 1b and c), we next wanted to determine whether or not PI4KIIIβ activates signaling pathways through modulation of membrane transport ^41^. To this end, we used chlorpromazine, an inhibitor of the formation of clathrin-coated pits, to halt clathrin-dependent endocytosis in BT549 cells. We observed that chlorpromazine decreases the activation of IGF-IRβ in wild type and ectopically expressing cell lines (Figures 8a and b). This non-toxic dose of chlorpromazine reduces endocytosis by 60-70% relative to controls (Supplementary Figure 3). Thus, endocytosis regulates IGF-IRβ activation and the increased activation observed in the PI4KIIIβ overexpressers (Figure 2c) depends on endosome function.

**Figure 8.**
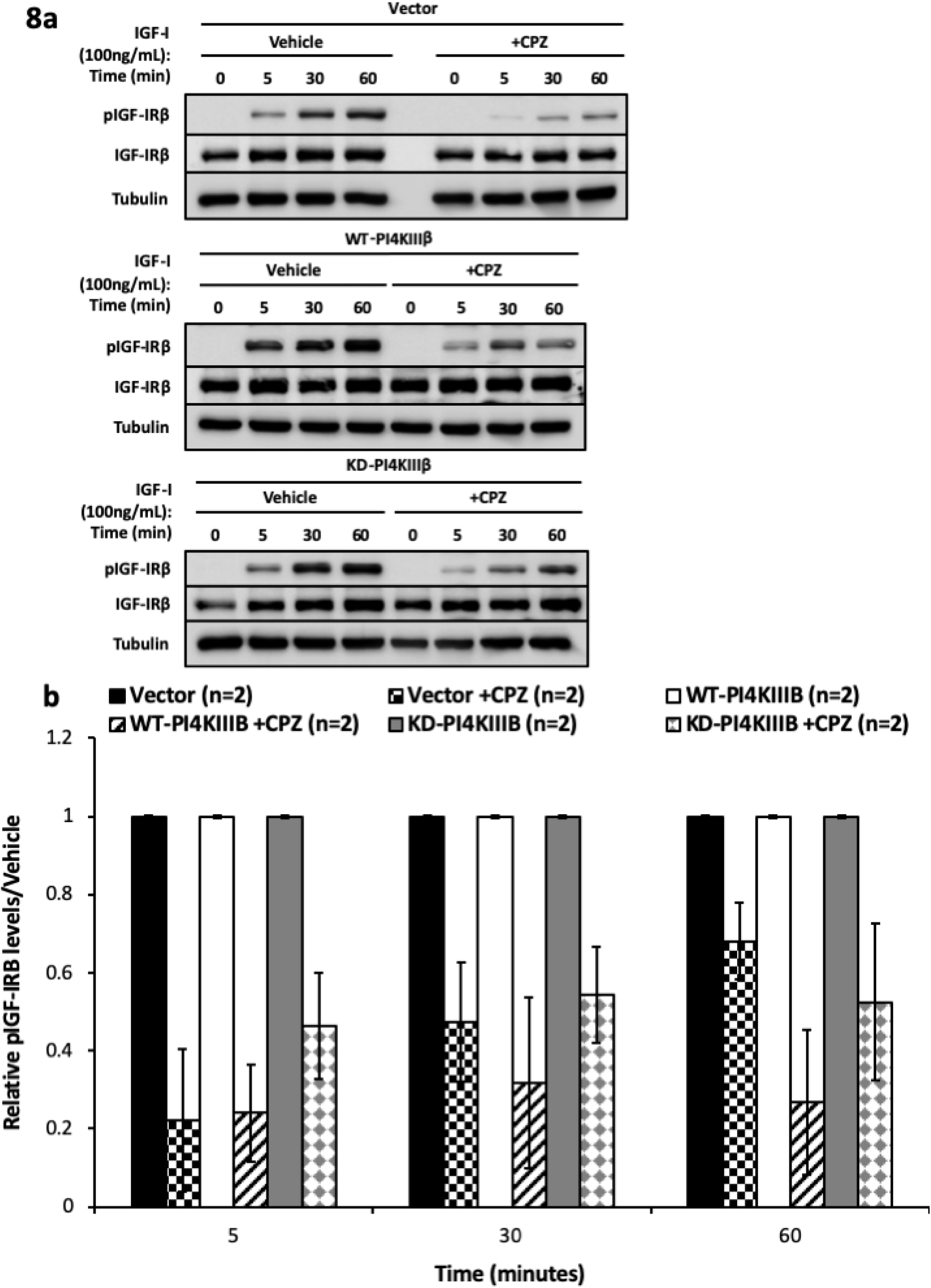
PI4KIIIβ regulates IGF-IRβ signaling through endosomes. (**a**) BT549 vector control (Vector), wildtype PI4KIIIβ-overexpressing (WT-PI4KIIIβ), and kinase-inactive PI4KIIIβ-expressing (KD-PI4KIIIβ) human breast ductal carcinoma cell lines were serum starved overnight followed by stimulation with IGF-I (100ng/mL) in the presence or absence of endocytic inhibitor, chlorpromazine (CPZ) (10μg/mL), for the indicated time periods. Lysate was collected and subjected to western blot analysis to determine levels of pIGF-IRβ (95kDa) and IGF-IRβ (95kDa) with Tubulin (55kDa) as a loading control. Each lane contains 30µg of total protein. (**b**) Protein levels were quantified by densitometry and the data shown represents the mean ± SE of the mean from 2 independent trials comparing the pIGF-IRβ levels relative to the IGF-IRβ levels in the Vector, WT-PI4KIIIβ, and KD-PI4KIIIβ cell lines in the presence of CPZ, followed by normalization to the same cell line with vehicle for each time point. Statistical significance (*, *P* < 0.05, one-way ANOVA with multiple comparison test) is indicated.

## DISCUSSION

In this study, we establish an important role for PI4KIIIβ in the regulation of the endocytic system. We show that PI4KIIIβ expression increases the rates of both internalization and recycling of transferrin. PI4KIIIβ deletion has the opposite effect, decreasing both internalization and recycling rates. Furthermore, loss of PI4KIIIβ decreases cellular Rab11a activity, suggesting that PI4KIIIβ-mediated control of endocytosis is mediated by regulating Rab11a function. Importantly, a kinase-inactive version of PI4KIIIβ rescues normal recycling in PI4KIIIβ-null cells, indicating that PI4KIIIβ-dependent control of recycling is independent of lipid kinase activity.

The PI4KIIIβ protein is highly expressed and the *PI4KB* gene amplified in a subset of primary human breast tumours^7, 28^. In addition, high PI4KIIIβ expression disrupts normal breast epithelial morphogenesis, resulting in irregular polarization and formation of multi-acinar structures ^31^. PI4KIIIβ has also been identified as a downstream effector of the known breast and ovarian cancer oncogene eEF1A2 and is involved in actin remodeling and cell motility^1, 19^. Here, we find that deletion of PI4KIIIβ decreases the rate of tumour development in mice. This observation is consistent with the idea that PI4KIIIβ is a breast cancer oncogene.

PI4KIIIβ-Rab11a interaction is required for PI4KIIIβ-mediated regulation of endosome function since a Rab11a binding mutant is unable to rescue recycling in PI4KIIIβ-null cells. Rab11a is necessary for the transport of recently internalized endosomes from the plasma membrane to the *trans*-Golgi network and the ‘slow’ recycling of internalized endosomes back to the plasma membrane^44–46^. Rab11a regulates these processes by directly binding effector proteins and recruiting them to endosomal vesicles. The PI4KIIIβ protein binds directly to Rab11a^5^ and has previously been shown to be required for the proper intracellular localization of Rab11^9, 32^. We observed that loss of PI4KIIIβ causes a drop in the overall Rab11a cellular activity as measured by Rab11 FRET biosensor. This is consistent with the idea that PI4KIIIβ is a direct activator of Rab11a. We hypothesize that PI4KIIIβ stabilizes interaction between GTP-loaded Rab11a and its effectors and thus promotes endosomal function. This potentiation of Rab11a activity would be independent of PI4KIIIβ lipid kinase activity. In cells with high levels of PI4KIIIβ, as would occur in approximately 20% of breast tumours, we hypothesize that increased number of PI4KIIIβ-Rab11a complexes increases the rate of endosome maturation, causing rapid delivery of internalized endosomes to the plasma membrane. Further work will be necessary to identify the Rab11a effectors involved.

Ectopic PI4KIIIβ expression not only increases the rate of the endocytic transport but also the activation of IGF-IRβ and Akt^28^. For IGF-IRβ, this activation is dependent on endosome function. We propose that PI4KIIIβ-mediated increase of endocytic rates amplifies the activation of plasma membrane signaling. Activation of of receptor tyrosine kinases (RTKs) and G-protein coupled receptors (GPCRs) is closely linked with the endocytic system^17, 45^. Following extracellular ligand-induced activation, the activated receptor complex enters the cell via clathrin-dependent endocytosis^12, 17^. Once internalized, activated receptor complexes may enter signaling endosomes^37, 38^. Here, the recruitment of adaptor proteins promotes the initiation of intracellular signaling events unique to endosomes^37, 38^. For example, in an early endosome containing activated EGFR, Rab5 will recruit APPL proteins to interact with Rab5 and EGFR; this will, in turn, promote the activation of Akt^24, 35^.

We propose that efficient endocytic recycling modulates signalling downstream IGF-IRβ. This is based on the finding that IGF-IRβ phosphorylation require PI4KIIIβ-mediated receptor recycling and a fully functional clathrin mediated endocytosis. Our model for PI4KIIIβ function is that in cancer cells with high PI4KIIIβ expression, activated receptor complexes are rapidly internalized and recycled back to the plasma membrane by trafficking into a PI4KIIIβ and Rab11a-positive endosome than in wildtype cells. Accordingly, increased number of PI4KIIIβ containing vesicles located near the periphery of the cell were measured. This process sustain the repeated activation of signaling events at the cell surface and in signaling endosomes. We propose that this endocytic-regulated control of signaling pathway activation contributes to the increased proliferation, survival and migration of those breast cancers with high PI4KIIIβ^42^.

Previous evidence has shown that the dysfunction of the endocytic system can contribute to the pathogenesis of cancer, through loose regulation of signaling events. For example, Rohatgi *et al* discovered a novel role for beclin 1 as a regulator of early endosome maturation^34^. They revealed that beclin 1, best characterized for its role in autophagy, regulates the maturation of early endosomes by recruiting phosphatidylinositol 3-phosphate (PI3P) in response to growth factor stimulation and that this control of early endosome maturation controls the duration of growth factor receptor signaling on endosomes^34^. The loss of *BECN1* in breast tumours, was shown to cause sustained activation of Akt and ERK in breast cancer cells^13^. Additionally, endosomal sorting complexes required for transport (ESCRT) complexes sort endosomes and, through post-internalization modifications, target the endosomes for recycling or degradation^36^. The loss of activity in subunits of ESCRT complexes is often found in human cancers and has been shown to be sufficient for the development of metastatic tumours^21, 22^. Elevated Rab11a expression has been observed in multiple cancers and can promote cell migration in cancerous cells of the gut, skin and breast ^14, 16, 18^. This suggests that Rab11a, and its activators and effectors, are likely to be oncogenes. However, this role may not be strictly universal since a recent report indicates that loss of Rab11a in mice promotes epithelial dysplasia of the intestine and reduced Rab11a expression is associated with poor survival in stage II and III colon cancers^8^.

We were surprised to see that PI4KIIIβ expression had an effect on endocytic internalization. Since Rab11a is not known to affect internalization, we believe that PI4KIIIβ is more extensively involved in the endocytic system than with just Rab11a and recycling endosomes. The Rab11a binding domain we disrupted in PI4KIIIβ (N162) interacts with Rab11a at a leucine residue that is conserved among other Rab proteins, including Rab5 and Rab7 ^5^. Rab5 controls the internalization of clathrin-coated pits^10^. Since we see a significant effect on transferrin internalization due to PI4KIIIβ expression, we hypothesize that PI4KIIIβ is acting with Rab5 to facilitate rapid internalization. This is consistent with our observation that PI4KIIIβ can be found in the same vesicles as Rab5. Additionally, termination of signaling is often achieved through the degradation of receptors and ligands via the late endocytic and lysosomal pathways ^38^. As Rab7, which we also observed to be found in the vesicles with PI4KIIIβ, is concentrated on late endosomes and is crucial for endo-lysosomal trafficking, it could possibly be involved or affected by PI4KIIIβ-mediated regulation of endocytic kinetics and signaling^10, 37^. Although PI4KIIIβ appears to have an important relationship with Rab11a, further work is necessary to determine how PI4KIIIβ and other endocytic regulatory proteins modify endocytic internalization. The importance of the PI4KIIIβ-Rab11a interaction in signaling activation suggests that this interface may be a druggable target for cancer therapy.

Long thought to simply be a way to move nutrients into and waste out of a cell, evidence is now emerging for the endocytic system as a regulator of cellular function and behaviour^25, 37, 38^. Studies implicate endocytosis in polarity, proliferation, migration, division, and transcription, among other cellular regulatory processes^11, 37, 38, 42, 47^. When appropriately regulated, this permits cells to survive in their environment and perform their intended functions. However, when regulation of the endocytic system is perturbed, this can lead to disruption in polarity, enhanced proliferation, abnormal migration, and uncontrolled division, ultimately leading to the development and progression of cancers^29, 37, 38^. Here, we reveal that PI4KIIIβ is a novel regulator of endocytic kinetics. Activation of endocytosis by PI4KIIIβ stimulates signaling activation and high expression of PI4KIIIβ in tumours results in the promotion of signaling processes that drive cancer initiation and progression.

## MATERIALS AND METHODS

### Cell lines and culture

The BT549 human breast ductal carcinoma cell line was obtained from ATCC (Manassas, VA, USA). The 4T1-luc mouse mammary carcinoma cell line constitutively expressing the firefly luciferase gene was a gift from Dr. John Bell (University of Ottawa, Canada). BT549 cells were cultured in RPMI-1640 from Life Technologies supplemented with 10% FBS (Thermo Scientific, Burlington, Canada), 1mmol/L sodium pyruvate (Thermo Scientific), 10nmol/L HEPES buffer (VWR International, Mississauga, Canada), 0.023 IU/mL insulin from bovine pancreas (Sigma-Aldrich, Oakville, Canada), and penicillin-streptomycin (Thermo Scientific). 4T1 cells were cultured in Dulbecco’s modified Eagle’s medium (Thermo Scientific) supplemented with 10% FBS, 1mmol/L sodium pyruvate, and penicillin-streptomycin. BT549 cell lines stably expressing ectopic PI4KIIIβ were created as described^6^. IGF-1 (catalog no. I3769) and chlorpromazine (catalog no. C8138) were purchased from Sigma-Aldrich. BT549 and 4T1 cell lines with PI4KIIIβ deletion by CRISPR/Cas9 were generated by transfecting wildtype cells with CRISPR/Cas9 plasmid (Santa Cruz Biotechnology, Mississauga, Canada) followed by single cell FACS of green fluorescent cells into 96-well plates. Single cells were grown to colonies and the deletion was verified by western blot and immunofluorescence. The PI4KIIIβ targeted RNA sequences for human and mouse were 5’-CCCTGATGGCGATCGGCAAG-3’, 5’-TCCTGCCAGCCGGCGCCTTT-3’, 5’-TATGAGCCAGCTGTTCCGAA-3’ (catalog no. sc-4185251) and 5’-CAGACCGTGTACTCCGAATT-3’, 5’-GGCTCCCTACCTGATCTACG-3’, 5’-ATAAGCTCCCTGCCCGAGTC-3’ (catalog no. sc-430739), respectively. Human cell lines have a deletion of exons 5-9 and mouse cells a deletion of exons 4-5. BT549 cell lines stably expressing rescued ectopic PI4KIIIβ were created as previously described except starting with BT549 PI4KIIIβ-null cell lines as the parental cells^6^.

### Western blot

Cells were lysed in radioimmunoprecipitation assay buffer (Tris-HCl, pH 7.4, 50mM; NaCl, 150mM; NP-40 1%; sodium deoxycholate, 0.5%; sodium dodecyl sulfate, 0.1%; ethylenediaminetetraacetic acid, 2mM; sodium fluoride, 50mM) supplemented with protease and phosphatase inhibitor cocktails (Roche, Mississauga, Canada). Protein concentrations were determined by Bradford protein assay (Bio-Rad, Mississauga, Canada). Loading buffer was added to 30μg of protein lysate and resolved by SDS-PAGE. The protein was then transferred onto a polyvinylidene difluoride membrane (Millipore, Toronto, Canada) and probed using antibodies for PI4KIIIβ (BD Biosciences 611817; Mississauga, Canada), vinculin (Santa Cruz Biotechnologies sc-25336), tubulin (Cell Signaling Technology 3873; Whitby, Canada), IGF-IRβ (Cell Signaling Technology 3027; Whitby, Canada) (catalog no. 3027) and phosphoIGF-IRβ Y1135/1136 (Cell Signaling Technology 3024; Whitby, Canada). Anti-mouse HRP-linked (catalog no. 7076), as well as anti-rabbit HRP-linked (catalog no. 7074) secondary antibodies were all obtained from Cell Signaling Technology. Bands were detected with a MicroChemi chemiluminescent system (DNR Bio-Imaging Systems, Toronto, Canada) and intensities were quantified by densitometry using GelQuant (DNR Bio-Imaging Systems).

### Immunofluorescence and flow cytometry

For immunofluorescence, BT549 cells were grown on 22 x 22mm #1.5 coverslips or ibidi (Madison, WI, USA) 35mm μ-Dishes for confocal experiments or World Precision Instruments (Sarasota, FL, USA) 35mm Fluorodishes for TIRF experiments and fixed with 3.7% paraformaldehyde in PHEM buffer for 10 minutes at 37°C. The cells were permeabilized with 0.5% Triton X-100 PBS for 10 minutes. The cells were blocked in Abdil (0.1% Triton X-100, 2% BSA-PBS) for 10 minutes. The cells were then incubated with primary antibodies in Abdil for 1 hour. Cells were washed 5 times with 0.1% Triton X-100 PBS and incubated with appropriate secondary antibodies in Abdil for 45 minutes. When dual staining, the previous steps were repeated from the blocking step with the necessary antibodies. The cells were stained with 1μg/mL DAPI for 5 minutes. The coverslips were then mounted on slides with Dako mounting medium and the dishes were mounted with ibidi mounting medium. The antibodies used for immunofluorescence staining were PI4KIIIβ (catalog no. AP8030a) from Abgent, (Mississauga, Canada), PI4KIIIβ (catalog no. 611817), EEA1 (catalog no. 610457), GM130 (catalog no. 610822) from BD Biosciences and Rab11a (catalog no. ab170134, ab3612), Rab5 (ab18211), Rab7 (ab50533), TGN46 (ab50595) from Abcam (Cambridge, MA, USA). The secondary antibodies used were all purchased from Invitrogen (Burlington, Canada) and include anti-mouse AlexaFluor®488 (catalog no. A-11029), anti-mouse AlexaFluor®546 (catalog no. A-11003), anti-rabbit AlexaFluor®488 (catalog no. A-11008), and anti-rabbit AlexaFluor®546 (catalog no. A-11010). All antibodies were used as per the manufacturer’s recommendations. Confocal images were acquired with a Zeiss LSM 510 META/AxioVert 200 confocal microscope with a 63x Plan-Apochromat 1.4 NA oil objective and Zen 2009 software. TIRF images were acquired with a TIRF-Spinning Disk Spectral Diskovery System with a 63x Plan-Apochromat 1.4 NA oil objective and MetaMorph software. All analysis of images was performed using ImageJ. Coincident fluorescence analysis was performed by measuring the plot profile of at least 50 vesicles in the periphery of the cells. Analysis of the TIRF images was performed by circling the outline of the cell and measuring the number, size, area, and circularity of the particles after background subtraction and thresholding.

For flow cytometry, cells were washed 2 twice in PBS and de-adhered with 5mM EDTA-PBS. 10^6^ cells in suspension were washed twice in PBS and resuspended in 100 uL of 0.2% BSA-PBS. 1 uL of anti-Transferrin Receptor Antibody (Abcam ab22391) was added and incubated for 30 minutes on ice. Cells were then washed twice in BSA-PBS and resuspended in 100 uL of BSA-PBS. 0.5 uL of anti-rat AlexaFluor®488 secondary antibody (Abcam ab 150165) was added for 30 minutes on ice. Cells were washed three times in BSA-PBS, resuspended to a volume of 300 uL and analyzed in a Beckman Coulter CyAN ADP9 using Summit v4.3.02 acquisition software. Files were analyzed using FlowJo software,

### Transferrin pulse-chase and uptake assays

BT549 cells were plated in 35mm μ-Dishes from ibidi to reach ∼70% confluency the following day. The cells were then washed with PBS twice and incubated in serum-free uptake medium (SFUM; RPMI-1640 with 10nmol/L HEPES buffer, penicillin, streptomycin and 0.1% BSA) for 1 hour at 37°C to deplete transferrin. For uptake experiments, cells were then incubated in SFUM containing 25μg/mL transferrin AlexaFluor®488 or 546 (Thermo Scientific) for 1 hour at 4°C (uptake assay). Cells were then transferred to 37°C for the indicated time. For recycling, after trasnferrin depletion, cells were incubated in SFUM containing 25μg/mL transferrin AlexaFluor®488 or 546 (Thermo Scientific) for 30 minutes at 37°C). Cells were then washed twice with SFUM to remove surface transferrin and incubated at 37°C with SFUM containing 1mg/mL unlabelled holo-transferrin (Sigma-Aldrich) (pulse-chase) or (uptake). At the specified times the cells were placed on ice and washed 3 times with PBS followed by a 6 minute wash with an acid wash buffer (20mM acetic acid, 500mM NaCl, pH 3.0). The cells were then fixed with 4% paraformaldehyde for 10 minutes at room temperature and permeabilized with 0.25% saponin, 1% BSA-PBS for 10 minutes on ice. Finally, the cells were incubated with 1μg/mL DAPI for 5 minutes and mounted with ibidi mounting medium. Images were acquired with a Zeiss LSM 510 META/AxioVert 200 confocal microscope with a 63x Plan-Apochromat 1.4 NA oil objective and Zen 2009 software. ImageJ was used to quantify the average corrected total cellular fluorescence (CTCF). For each time point and cell line at least 25 cells were outlined and the total integrated density was determined. CTCF was determined as previously described by multiplying the area of selected cell by the mean fluorescence of 4 background readings and subtracting that from the integrated density (CTCF = integrated density – (area of selected cell x mean fluorescence of background readings)^4^. The CTCF values were then normalized to the 0 time point in order to account for any differences in transferrin receptor expression between the cell lines. For the pulse-chase assays, the CTCF values were normalized to the average CTCF of the 0 time point and inverted in order to determine the relative transferrin recycled (decrease in cellular fluorescence). For the uptake assays, the CTCF values then had the average CTCF for the negative control subtracted and were normalized to the average CTCF of the binding control in order to determine the relative transferrin internalized (increase in cellular fluorescence).

### *In vivo* tumour studies

Animals were housed and handled according to Canadian Council on Animal Care standards and policies. 100,000 4T1-luc cells/mouse in PBS were injected into the mammary fat pads of 7-9 week old Balb/c mice. For imaging, mice were injected *i.p.* with 200μL of 15mg/mL D-Luciferin (PerkinElmer, Waltham, MA, USA) in PBS and luminescence quantitated 5 minutes later using a PerkinElmer IVIS SpectrumCT.

## ACKNOWLEDGEMENTS

The authors thank Mohsen Alipour for assistance with proximity ligation assays and Durga Sivanesan and Catherine St-Louis for technical assistance. The authors also thank Skye McBride, Chloe van Oostnde, Tong Zhang, Colton Boudreau and Erica Tse-Luen for training and assistance with microscopy and Vera Tang for help with the flow cytometry. The PI4KIIIβ-N162A was a gift from Dr. John Burke. We thank John Copeland, Jim Dimitroulakos, Denise Law-Vinh and Robyn Skillings for helpful discussion and critical reading of this manuscript. This work is supported by operating grants from the Canadian Cancer Society Research Institute (JML and RA).

## SUPPLEMENTARY INFORMATION

**Figure S1.** PI4KIIIβ does not affect transferrin receptor expression. Flow cytometry quantitation of transferrin receptor (CD71) expression in BT549 vector control (Vector), wildtype PI4KIIIβ-overexpressing (WT-PI4KIIIβ), and kinase-inactive PI4KIIIβ-expressing (KD-PI4KIIIβ) human breast ductal carcinoma cell lines. Inset table shows expression means in three independent experiments. Differences between Vector and either WT-PI4KIIIβ or KD-PI4KIIIβ lines are not significant (t-test, p<0.57 and p<0.40 respectively)

**Figure S2.** PI4KIIIβ does not substantially affect Rab11a recycling endosome size or number per cell, number, shape, or size. *Upper panel.* The size of Rab11a vesicles in BT549 wildtype (Wildtype, n=13), two independent CRISPR/Cas9 knockouts of PI4KIIIβ (null A and B, n=26 and n=24), vector control (Vector, n=18), wildtype PI4KIIIβ-overexpressing (WT-PI4KIIIβ, n=16), kinase-inactive PI4KIIIβ-expressing (KD-PI4KIIIβ, n=15), and cells transfected with active Rab11a (Rab11a, S20V). Data is presented as the mean ± SD relative to Wildtype cells. Statistical significance (*, *P* ≤ 0.0001, t-test) is indicated. The number of Rab11a vesicles per cell area is presented as the mean relative to Wildtype cells ± SD. Statistical significance (*, *P* ≤ 0.001, t-test) is indicated. Cells transfected with constitutively active Rab11a (S20V) show larger but fewer Rab11a vesicles. *Lower panel.* Representative images of Rab11a vesicles in the cells.

**Figure S3.** Chlorpromazine inhibits endocytosis in BT549 cells. (**a**) Treatment of BT549 vector control (Vector), wildtype PI4KIIIβ-overexpressing (WT-PI4KIIIβ), and kinase-inactive PI4KIIIβ-expressing (KD-PI4KIIIβ) human breast ductal carcinoma cell lines with chlorpromazine or Dynasore reduces transferrin uptake after 30 minutes at 37°C. Statistical significance (*, *P* < 0.05, one-way ANOVA with multiple comparison test) is indicated. (**b**) Representative image from (**a**) showing wildtype cells. Scale bar is 50μM.

